# Human adult neurogenesis loss underlies cognitive decline during epilepsy progression

**DOI:** 10.1101/2022.09.12.507339

**Authors:** Aswathy Ammothumkandy, Luis Corona, Kristine Ravina, Victoria Wolseley, Jeremy Nelson, Nadiya Atai, Aidin Abedi, Nora Jimenez, Michelle Armacost, Lina M. D’Orazio, Virginia Zuverza-Chavarria, Alisha Cayce, Carol McCleary, George Nune, Laura Kalayjian, Darrin Lee, Brian Lee, Christianne Heck, Robert H. Chow, Jonathan J Russin, Charles Y. Liu, Jason A.D. Smith, Michael A. Bonaguidi

## Abstract

Mesial temporal lobe epilepsy (MTLE) is a syndromic disorder presenting with seizures and cognitive comorbidities. While seizure etiology is increasingly understood, the pathophysiological mechanisms contributing to cognitive decline and epilepsy progression remain less recognized. We have previously shown that adult hippocampal neurogenesis, a process contributing to visual spatial learning and memory in rodents, dramatically declines in MTLE patients with increased disease duration. Here, we investigate when multiple cognitive domains become effected by epilepsy disease duration and how human neurogenesis levels contribute to it. We find that intelligence, and verbal learning and memory decline at a critical period of 20 years disease duration. Surprisingly, the number of human immature neurons positively associate with auditory verbal, rather than visuospatial, learning and memory. Moreover, we uncover cognitive functions enriched to either immature or mature granule neurons and functions shared between them. Our study provides cellular evidence of how adult neurogenesis contributes to human cognition, and signifies an opportunity to advance regenerative medicine for patients with MTLE and other cognitive disorders.

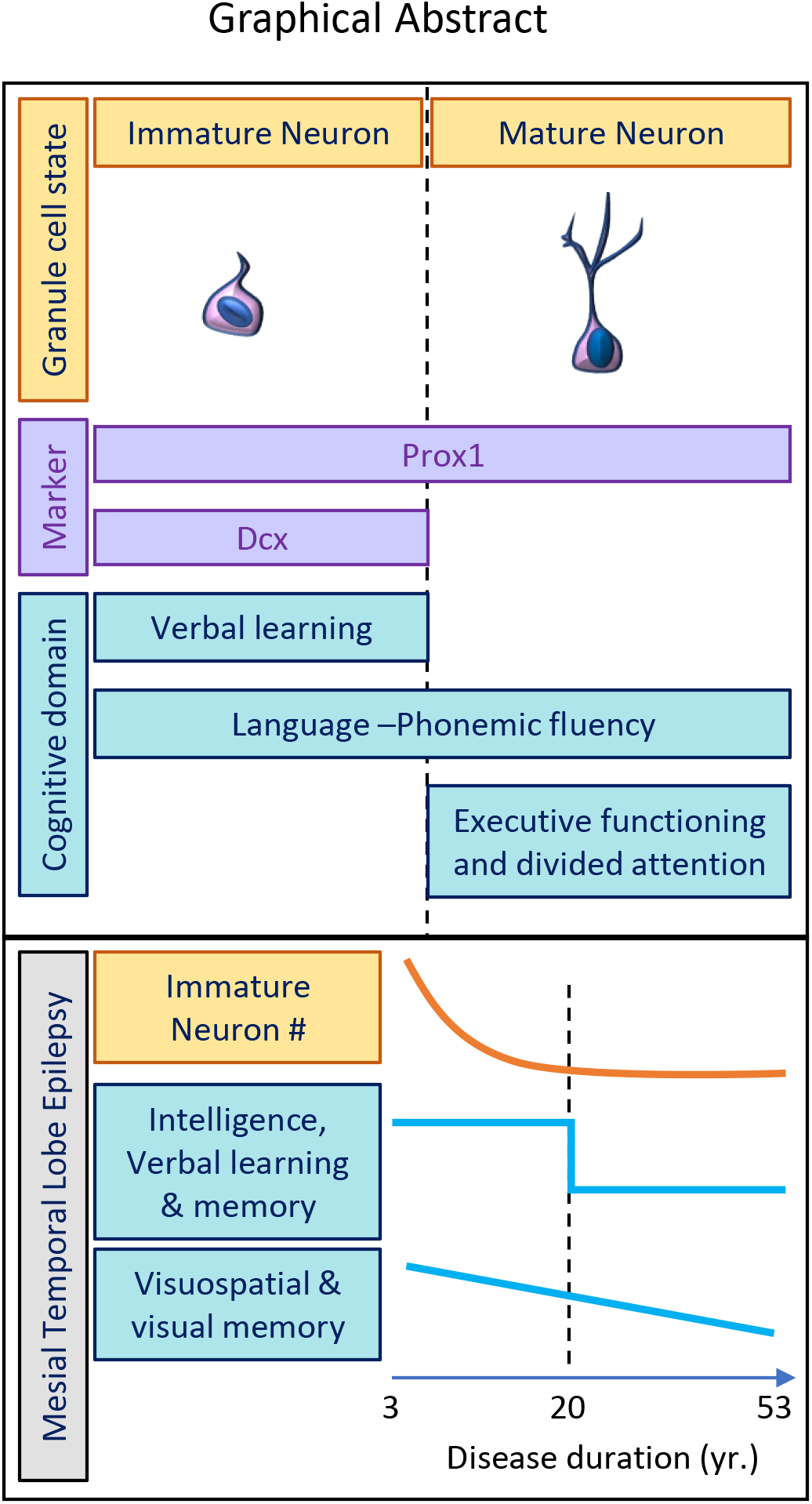

## INTRODUCTION

Epilepsy is a prevalent neurological disorder defined by recurrent seizures and presents comorbidities including neurocognitive dysfunction. Mesial temporal lobe epilepsy (MTLE) is the most common and refractory adult epilepsy, with 70% of patients becoming pharmacoresistant^1^. In comparison to other epilepsies, MTLE patients are more likely to exhibit disease progression including worsening seizure control, structural abnormalities, cognitive decline, and behavior impairment^2^. Cognitive phenotypes in MTLE have been categorized according to clinical criteria and data driven cluster analyses^3^. Approximately half of the patients possess intact cognition, while roughly one quarter display impairments in learning, memory and language; and another quarter display more widespread cognitive dysfunction including impaired attention, executive function and processing speed^3–5^. However, when and which structural abnormalities contribute to potential progressive cognitive impairment are not well understood in epilepsy patients.

MTLE is characterized by focal seizures in the hippocampus, which performs episodic learning and memory, spatial cognition, and emotional processing functions^6, 7^. The human hippocampus also contains a rare niche in which adult neurogenesis can persist throughout life^8–13^. In this process, neural stem cells produce proliferating neuronal progenitors that differentiate and mature into granule neurons. Functionally, adult immature neurons benefit visuospatial cognition and impede memory interference in rodent models of aging and dementia^14–17^. However, their role in human cognition remains unknown. A mild decline in hippocampal neurogenesis occurs in healthy people with age^12^, which is further exacerbated in Mild Cognitive Impairment (MCI) and Alzheimer’s disease (AD)^11, 13^. Our recent study^18^ on human MTLE shows the most profound decline in neurogenesis among patients with neurological diseases to date^11, 12, 19^. Neurogenesis rapidly deteriorates over the course of the disease and immature neurons were not detected after 20 years of MTLE disease duration (DD)^18^. This dynamic range provides an opportunity to define the functional role of neurogenesis in human cognition.

Prior studies indicate that MTLE patients with severe cognitive decline have longer mean DD^3, 20^ and implicate that cognitive impairment could be progressive^3, 20, 21^. Here, we examine a cohort of Spanish-speaking MTLE patients to identify which cognitive domains are affected by DD, and when they significantly decline. Sequential neuropsychological and histological analysis in the same MTLE patients reveal which cognitive domains are associated with the number of immature and mature dentate granule neurons. Our results link the decline and eventual loss of immature neurons to verbal learning impairment during a critical period of epilepsy progression.

We also observe impacted cognitive domains that are shared or distinct between immature and mature adult granule neurons.

## RESULTS

### Cognitive performance declines with advancing disease duration

Cognitive functioning was evaluated using the Neuropsychological Screening Battery for Hispanics (NeSBHIS)^22, 23^. We first identified which cognitive domains from the NeSBHIS (Table S1) were affected by DD. Education-normalized NeSBHIS z-scores and age plus education normalized z-scores for Grooved Pegboard fine motor test^22^ were analyzed from MTLE patients (N = 36), average age (M = 39.83 years; SD = 11.54), DD (M = 27.43 years; SD = 13.77) and reported years of education (M = 8.03 years; SD = 3.51) (Table S2). Spearman’s correlation revealed declining performances across multiple cognitive domains (Table S3). Hippocampal-dependent functions of verbal learning and memory^24^ significantly declined on learning (Figure 1b), and short delay recall trials (Figure 1c) tests. The verbal long delay recall trial (Figure 1d) also declined with DD, trending toward statistical significance. In addition to verbal domains, reasoning and intelligence (Figure 1a)^20^, visuospatial skills and visual memory (Figure 1e-f) showed significant declines with DD. The remaining cognitive domains including language and executive functioning did not show significant associations with DD (Figure S1). Hence, this cohort of Spanish-speaking MTLE patients present progressive cognitive decline with DD in intelligence and both verbal and visuospatial memory domains.

**Figure 1.**
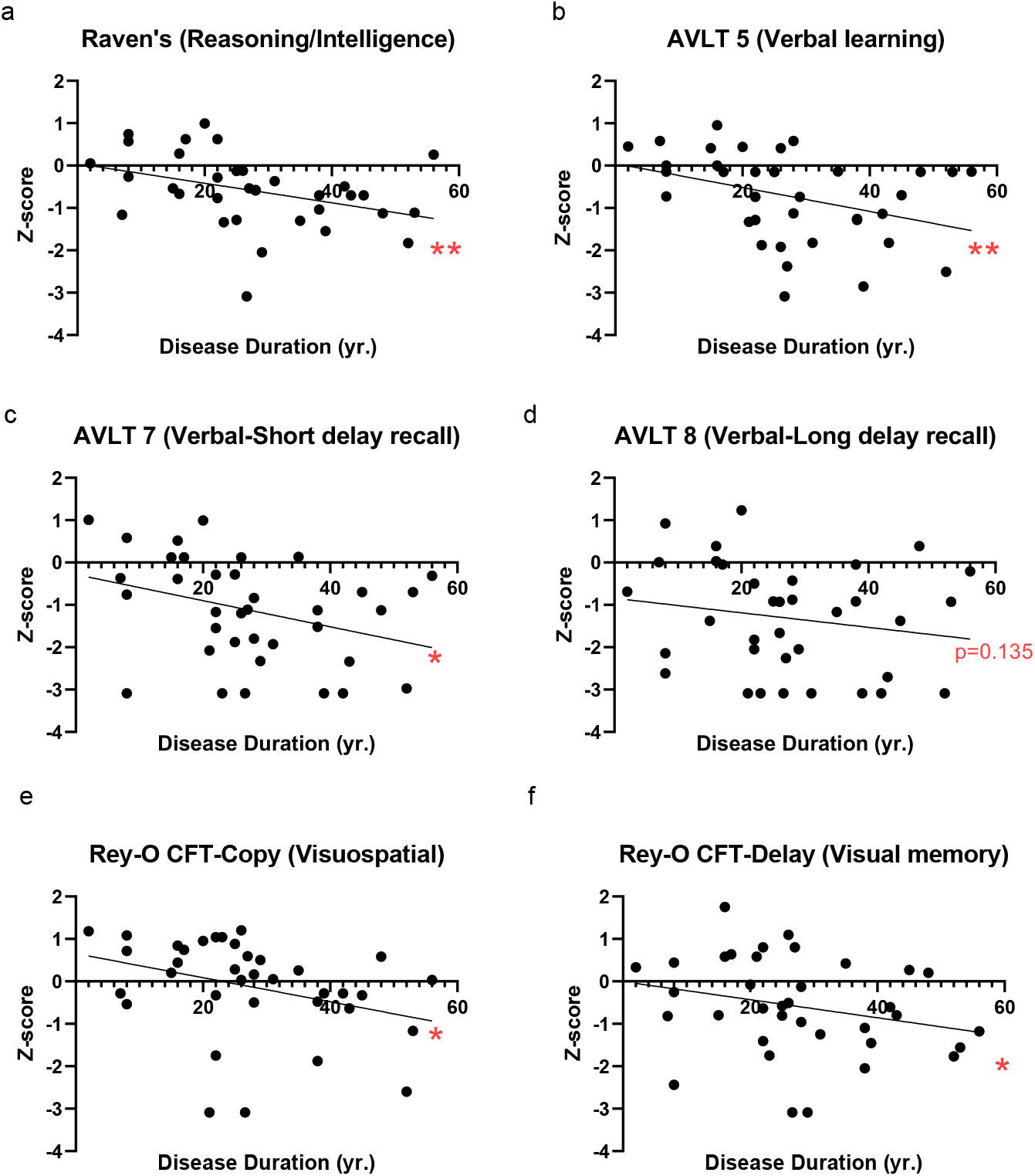
Cognitive performance domains declining with advancing disease duration (DD). Association between DD and cognitive z-scores calculated by two-tailed Spearman’s correlation for NeSBHIS tests: a) Raven’s Intelligence, b) AVLT 5 verbal learning, c) AVLT 7 verbal short- delay recall, d) AVLT 8 verbal long-delay recall, e) Rey-O-CFT Copy visuospatial, f) Rey-O-CFT Delay visual memory, * P ≤ 0.05, ** P ≤ 0.01.

### A critical period of cognitive impairment in MTLE patients

To identify the critical period when cognitive performance significantly decreases, we separated the patient data into three distinct groups to have approximately similar n-value based on DD. Group 1 (G1, N = 10) was < 20 years DD, group 2 (G2, N = 14) was 21 to 30 years DD, and group 3 (G3, N = 12) contained > 31-years DD. We then compared whether the NeSBHIS z-scores differed significantly between groups (Table S4). With respect to intelligence (Figure 2a) and verbal learning and verbal short delay recall memory (Figure 2b-c), G2 and G3 had significantly lower z-scores compared to G1; meanwhile, scores did not decline significantly from G2 to G3. Verbal long delay recall (Figure 2d) also showed a similar trend with significant or close to significant lower z scores for G2 and G3 compared to G1, with not much difference between G2 and G3. This result indicates a sudden drop in performance rather than a slight gradual decline that was observed in visuospatial skills and visual memory (Figure 2e, f). The fine motor test for the dominant and non-dominant hand also displayed a significant drop in performance between G1 and G2 (Figure S2h, i). The remaining tests did not display significant differences between any DD groups (Figure 2 e, f, Figure S2a-g). Therefore, the most significant declines in intelligence, verbal learning and memory, but not visuospatial skills and memory, occur in MTLE patients before a critical period of 20 years (Figure 2g); revealing a window for therapeutic interventions to reduce the impact of specific cognitive comorbidities.

**Figure 2.**
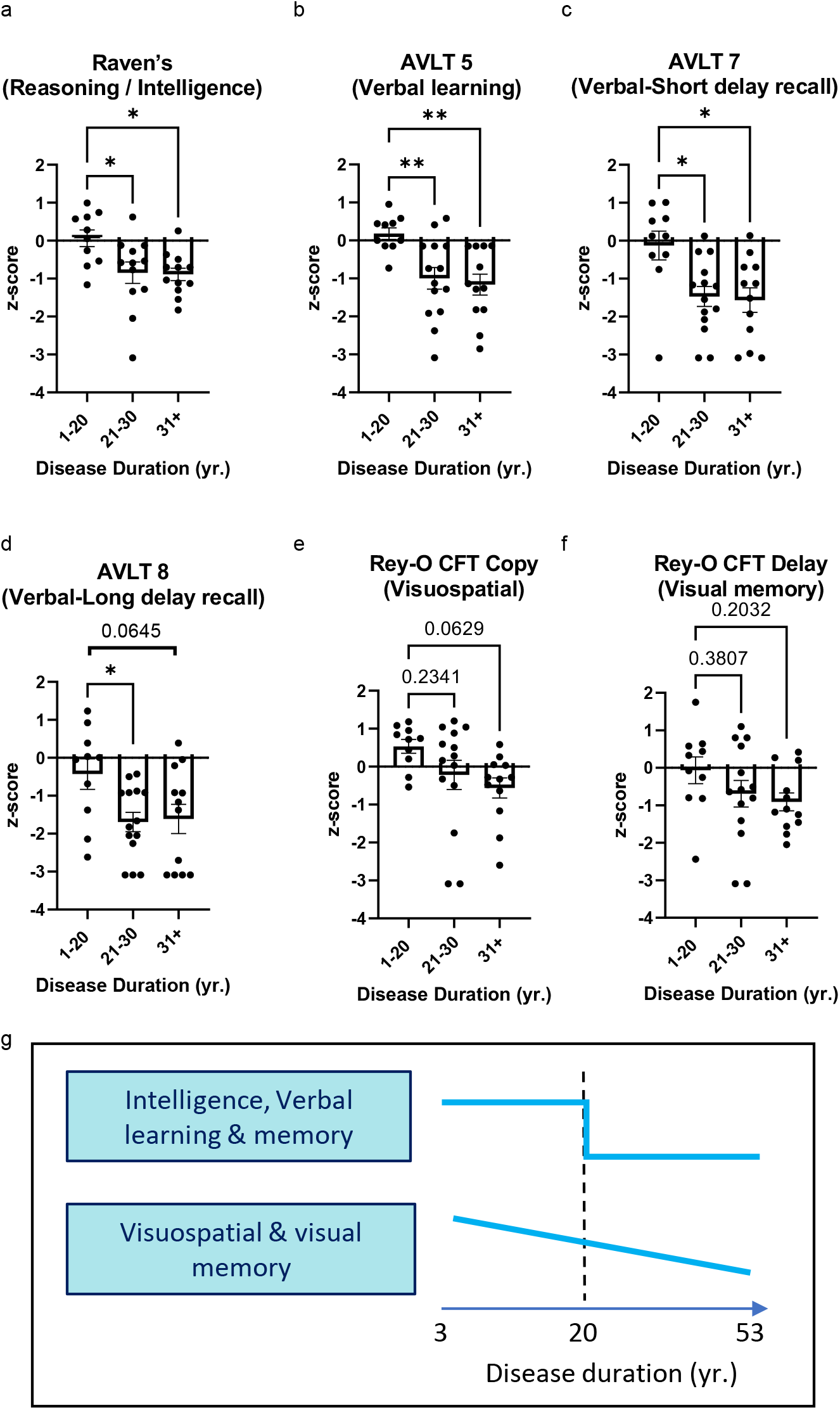
**A Critical period of cognitive impairment in MTLE patients**. Three DD groups were created; G1: 1-20 years DD, G2: 21-30 years DD, and G3: 31+ years DD. Cognitive z- scores between 3 groups were compared with a one-way ANOVA with post-hoc Tukey’s analysis: a-d) Cognitive z scores were significantly or close to significantly higher for G1 compared to G2 and G3 for: a) Raven’s intelligence, b) AVLT 5 verbal learning, c) AVLT 7 verbal short-delay recall, d) AVLT 8 verbal long-delay recall; e, f) Cognitive z scores were not significantly different between groups. e) Rey-O CFT Copy visuospatial f) Rey-O CFT Delay visuospatial memory. * P ≤ 0.05, ** P ≤ 0.01. All error bars represent S.E.M across individual cases. g) Intelligence, verbal learning and memory decline at an early critical period, whereas visuospatial skills and memory decline progressively.

### Contributions of immature and mature granule neurons to cognition

The identified critical period (Figure 2) coincides with the disease progression when immature neurons are reported to become nearly undetectable in MTLE patients^18^. We next determined the relationship between the loss of immature neurons and cognitive performance in intelligence, verbal learning, and memory tests. We first benchmarked our previous observation of immature neuron decline with DD to our current cohort of Spanish-speaking patients only (the current cohort contains a subset of patients from our previous study with 4 additional cases). Immature neurons were quantified as the number of Dcx+Prox1+ cells (Figure 3a-b) by performing histology in surgically resected tissue (N = 17). Consistent with our previous report^18^, we observed an exponential decline of adult immature neuron numbers with DD (Figure 3c). We then analyzed the number of mature granule neurons (Prox1+ cells) with DD and observed a mild non-significant level of neurodegeneration (Figure 3d). Next, we investigated the association of immature and mature granule neuron levels across various cognitive domains (Table S5). We observed that verbal learning and language (phonemic fluency) showed a significant positive association with immature neuron levels (Figure 3 e, g). To further unravel whether this association is enriched to immature granule neurons, we investigated the contribution of mature granule neurons to various cognitive domains. Mature granule neuron levels showed no significant association with verbal learning (Figure 3f), but correlated positively with phonemic fluency (Figure 3h). Therefore immature, rather than mature granule neuron levels particularly benefit verbal learning; whereas both immature and mature granule neurons contribute to phonemic fluency (language). Meanwhile, executive function and divided attention positively correlated with the number of mature granule neurons rather than immature neurons (Figure 3 i, j). Neither immature nor mature granule neurons levels displayed a significant association with any other examined cognitive domains (Figures S3-4). Hence, our study identifies that immature and mature granule neuron numbers can be associated with distinct cognitive functions (Figure 3k) and provides cellular evidence for a specific contribution of neurogenesis stages to human cognitive performance.

**Figure 3:**
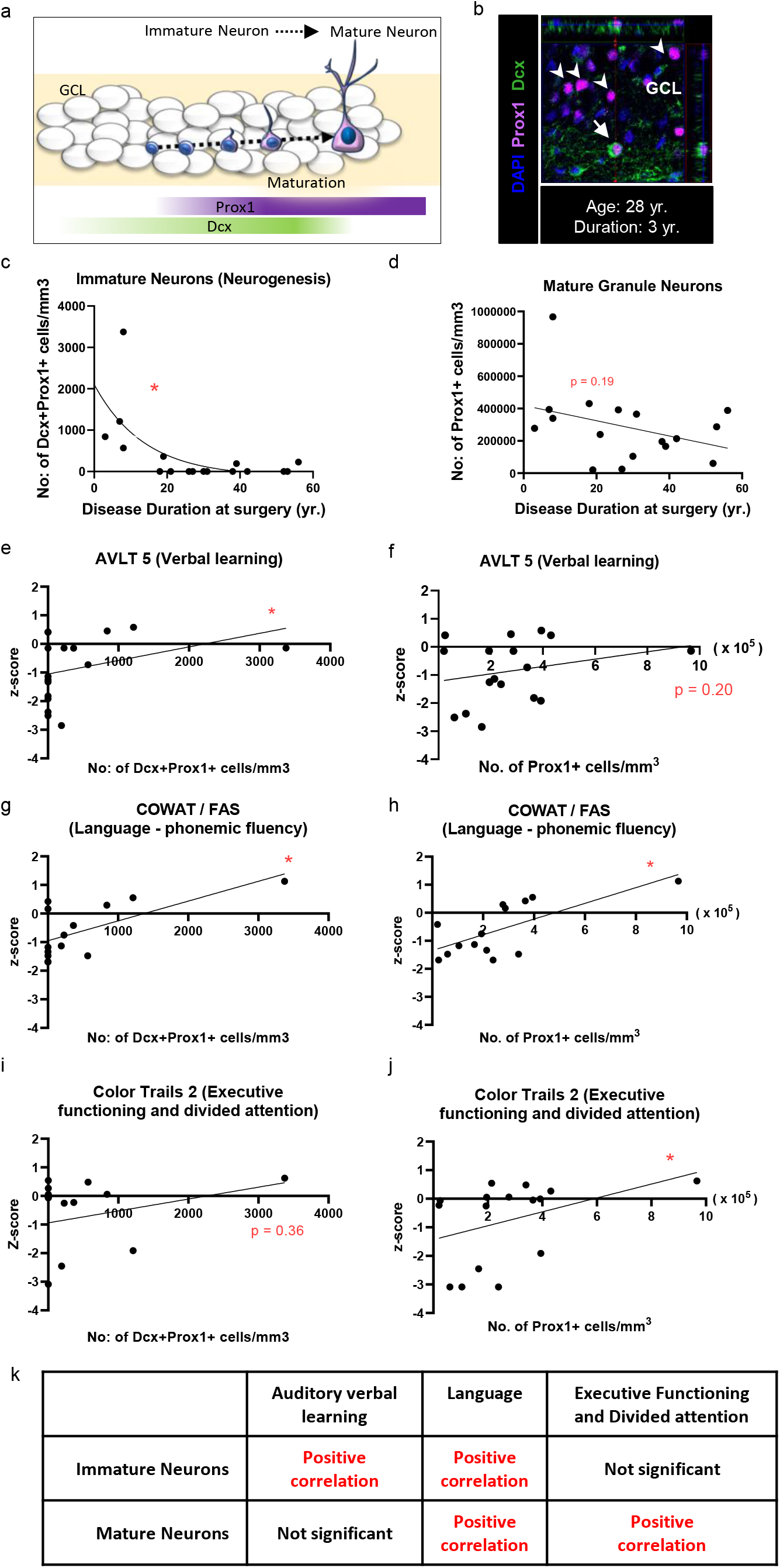
Contributions of immature and mature granule neurons to cognition. a) Cartoon illustration of Dcx and Prox1 expression across stages of granule neuron maturation. b) Dcx+ (green) Prox1+ (purple) immature neurons (Arrow) and Dcx- Prox1+ mature granule neurons (Arrowhead) identified in the Granular Cell Layer (GCL) of adult human MTLE cases. c) Number of Dcx+ Prox1+ immature neurons/mm3 in the GCL with DD (yr.) at surgery. Two-tailed Spearman’s Correlation * P ≤ 0.05. Spearman’s r = -0.5711. d) Number of Prox1+ granule neurons/mm3 in the GCL with DD (yr.) at surgery. Two-tailed Spearman’s Correlation P = 0.19 Spearman’s r = -0.3311. e, g, i) Two-tailed Spearman’s Correlation between the number of Dcx+ Prox1+ immature neurons/mm3 detected from the surgically resected hippocampus and presurgical cognitive z-scores for e) AVLT 5 verbal learning, g) COWAT/FAS language fluency, i) Color trails 2 executive function, * P ≤ 0.05. e, g. Two-tailed Spearman’s Correlation between number of Prox1+ granule neurons/mm3 detected from the surgically resected hippocampus and presurgical cognitive z scores: f) AVLT 5 verbal learning, h) COWAT/FAS language fluency, j) Color trails 2 executive function. * P ≤ 0.05. k) Summary table showing the significant correlations between the number of immature and mature granule neurons across cognitive domains.

## DISCUSSION

In MTLE, when and which structural abnormalities contribute to the progression of cognitive impairment are not well understood. Here, we find that human adult immature neuron loss underlies auditory verbal learning (AVLT5) impairment during a critical disease duration of 20 years. Although we observed a similarly pronounced initial decline in intelligence (Raven’s) and verbal memory (AVLT 7, -8) plus a gradual decline in visuospatial skills (Rey-O Copy) and visual memory (Rey-O Delay), these domains are less associated with immature neuron levels. In addition, we detected a link between executive function (Color Trails 2) and the number of mature granule neurons rather than immature neurons, while language fluency (FAS) is shared between the two granule cell states. These observations illustrate an interplay between neurogenesis levels and the progression of cognitive decline across multiple domains in MTLE patients (Graphical Abstract).

The role of immature neurons in specific human cognitive functions has remained unknown. A prior study of MTLE patients related lower proliferation of neural progenitors *in vitro* to impaired memory on the intracarotid sodium amobarbital test^25^. More recently, findings in patients with MCI identified that proliferating neuroblast numbers benefit global cognition^13^. Our analysis of multiple cognitive domains was critical in understanding the specific neuropsychological functions influenced by adult human neurogenesis. Remarkably, findings that levels of immature neurons, more so than mature granule neurons, particularly benefit verbal learning in MTLE patients alludes to a specialized function of adult neurogenesis. In rodents, new born granule cells possess enhanced intrinsic excitability, stronger responsiveness to external inputs^26^ and enhanced synaptic plasticity^7, 27, 28^, which contributes to learning, working memory and memory consolidation. Whether these properties are evolutionarily conserved in human new born granule neurons remains to be tested. Meanwhile, we find that the numbers of mature granule neurons, rather than immature neurons, positively associate with executive function and divided attention. Executive function is ascribed to the prefrontal cortex and its coupling with the temporal lobe^29, 30^. Our results therefore suggest that cognitive flexibility in MTLE patients could become impaired through mature granule neuron loss and altered functional connectivity between the temporal and frontal lobes^31^. These findings contrast with another study that links granule neuron loss with learning impairment in epilepsy patients^32^. Such divergent interpretations may be due to differences in the extent of granule cell loss and the learning and memory tests performed. Indeed, mature granule neurons facilitate memory recall^7^, which we also observe via the FAS language test that involves rapid retrieval of long-term crystalized, verbal knowledge of language, or semantic memory. The shared benefit of immature and mature neurons to outcomes on the FAS language test might reflect cumulative granule neuron turnover - neuronal cell loss and cell genesis - during human lifespan^8^. Thus, the current neurogenesis findings provide a new understanding of the cellular dynamics that link dentate gyrus volume change to cognitive outcomes^24, 33^.

Learning, memory, and executive function impairment reflect hippocampal network abnormalities that extend beyond epilepsy to other neurodegenerative disorders^6, 34, 35^. As individuals age, learning and working memory diminishes^36^, and is often accompanied in MCI by the beginning of executive dysfunction^37^. Early stage working memory and executive dysfunction in MCI can also predict progression to AD^35, 37–40^. Further, divided attention and episodic memory deficits begin to emerge with the onset of AD ^41^. Hippocampal neurogenesis deficits have been reported in such patients at early stages of both MCI and AD^11, 13^. The current study reveals impacted cognitive domains that are shared or distinct between immature and mature adult granule neurons. These findings serve as a crucial basis for identifying potential clinical interventions aimed at preventing or reversing cognitive decline in MTLE and other cognitive disorders. For example, a previous study in healthy individuals has also implied a role for exercise-induced neurogenesis on verbal learning using indirect measures of dentate gyrus cerebral blood volume^42^. Our more direct evidence quantifying the amount of immature and mature granule neurons in hippocampal resections from MTLE patients indicates neurogenesis benefits multiple cognitive domains. Further research is needed to understand whether therapeutic or lifestyle changes that promote neurogenesis and cognition in model systems can be as regenerative in humans.

### Study limitations

This study is moderately sized, which limits our ability to detect less dramatic cell type associations with cognitive domains. We are also insufficiently powered to analyze the contribution of neurogenesis lateralization between left and right hippocampus to verbal and visual memory^43^.

Finally, we do not conclude that neurogenesis lacks contribution to visuospatial learning and memory in people as has been related in rodent studies^16^. Such species divergence could reflect our study not assessing repeated visual learning and encoding. Future studies may investigate visuospatial learning using encoding tests such as the Biber Figure Learning Test (BFLT) that requires learning a series of figures presented over repeated trials^44^ or visual recognition memory performance (pattern separation) using the mnemonic similarity test (MST)^45, 47^.

## Author contributions

A.AK. and M.A.B. conceived the project. A.AK, L.C, R.H.C, D.S, C.Y.L, J.J.R, J.A.D.S, and M.A.B designed the experiments. A.AK, L.C, K.R, V.W, and J.N performed the experiments. N.J, L.M.D, V.C, C.M, K.R, N.A, A.A and A.AK compiled the clinical information. J.J.R, C.Y.L. D.L. and B.L performed the neurosurgeries. G.N, L.K, and C.H conducted clinical review and assisted with specimen collection. L.C and A.AK analyzed and compiled the data. L.C, A.AK, J.A.D.S and M.A.B wrote the manuscript. C.Y., J.A.D.S and M.A.B supervised the project.

## Data and materials availability

All data are available in the manuscript or supplementary materials. Any information regarding the availability of raw data, materials and methods can be directed to M.A.B.

## SUPPLEMENTAL INFORMATION

### Extended Figure Legends

**Table S1:** Cognitive domains and the respective NeSBHIS (Neuropsychological Screening Battery for Hispanics) tests.

| Cognitive Domain | Test (Description) |
| --- | --- |
| **Reasoning / Intelligence** | Raven's Standard Progressive Matrices<br>(Nonverbal/visual reasoning task to estimate general intelligence. Often regarded as culture fair measure.) |
| **Attention, Processing Speed and Executive Functioning** | Digit Symbol (Processing speed and graphomotor speed)<br>Color Trails 1 (Visual attention and processing speed)<br>Color-Trail 2 (Divided attention, cognitive flexibility, executive functioning) |
| **Language / Verbal Ability** | Pontón-Satz Boston Naming Test (P-S BNT; confrontational object naming drawn figures)<br>Controlled Oral Word Association Test (COWAT, FAS; measure of phonemic verbal fluency)<br>Digit Span (Verbal Attention and working memory, repetition of digits forward and then reverse) |
| **Verbal Learning and Memory** | World Health Organization – University of California, Los Angeles Auditory Verbal Learning Test (WHO-UCLA AVLT).<br>Trials 1-5 (Fifteen words presented over five trials to promote learning)<br>Trial 6 (Second 15-word list, an interference trial)<br>Trial 7 (Short delay recall of the original items, immediately follows Trial 6)<br>Trial 8 (20-minute, long delay recall) |
| **Visuospatial** | Block Design (Visuospatial constructional ability)<br>Rey-Osterrieth Complex Figure Test (ROCFT) - Copy (Reconstruction of an intricate, two-dimensional geometric figure) |
| **Visual Memory** | ROCFT – Delay (10-minute delayed recall, reconstructing the ROCFT stimuli) |
| **Fine Motor Functioning** | Grooved Pegboard Dominant (GP-D) and Nondominant (GP-ND) hands<br>(task of manual dexterity, visuomotor integration and speed) |

**Table S2:**
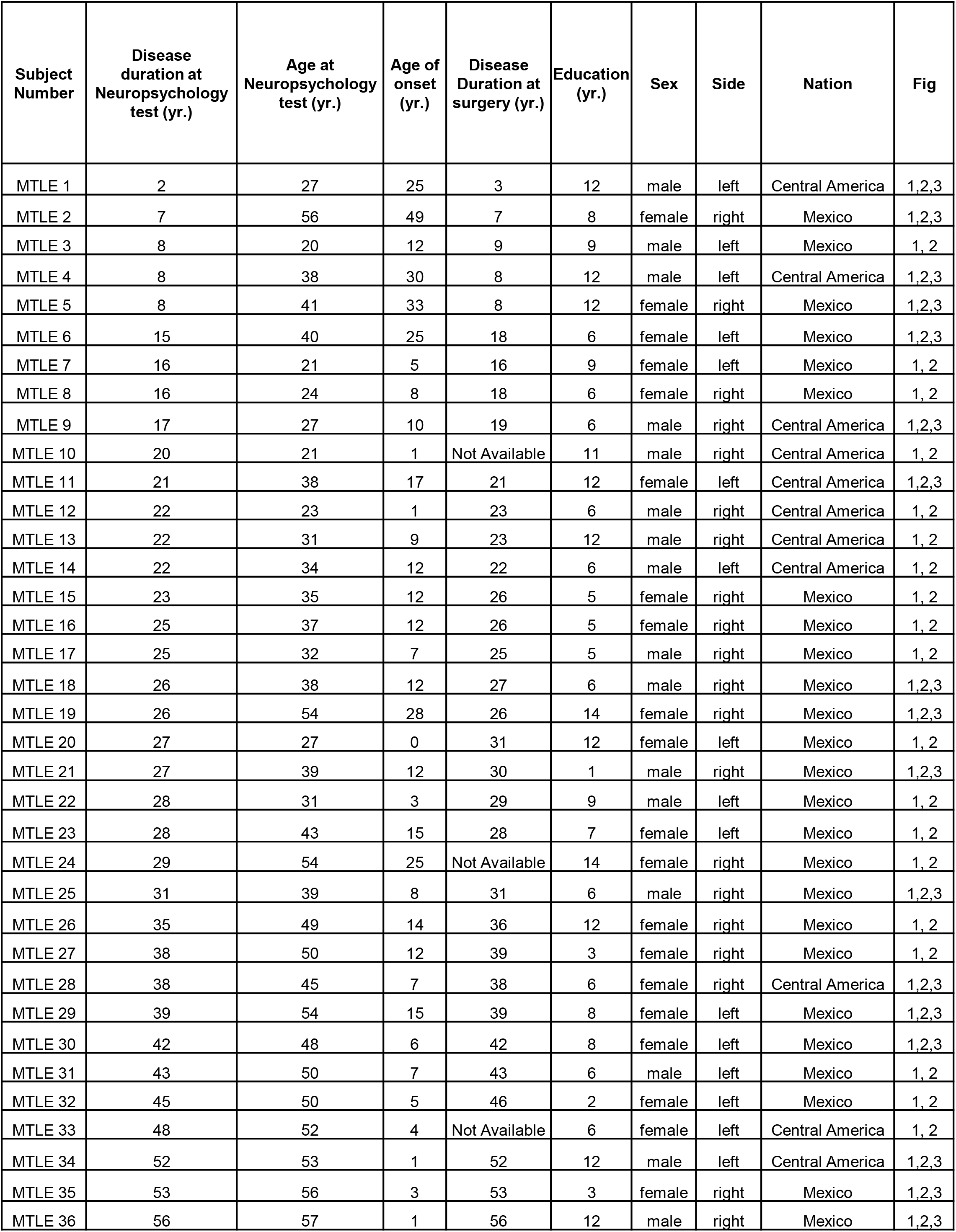
Clinical, demographic information and experiments performed in MTLE cases (N = 36). MTLE- Mesial Temporal Lobe Epilepsy Disease duration, Age at surgery, Age of onset and Education in years (yr.)

| Subject Number | Disease duration at Neuropsychology test (yr.) | Age at Neuropsychology test (yr.) | Age of onset (yr.) | Disease Duration at surgery (yr.) | Education (yr.) | Sex | Side | Nation | Fig |
| --- | --- | --- | --- | --- | --- | --- | --- | --- | --- |
| MTLE 1 | 2 | 27 | 25 | 3 | 12 | male | left | Central America | 1,2,3 |
| MTLE 2 | 7 | 56 | 49 | 7 | 8 | female | right | Mexico | 1,2,3 |
| MTLE 3 | 8 | 20 | 12 | 9 | 9 | male | left | Mexico | 1, 2 |
| MTLE 4 | 8 | 38 | 30 | 8 | 12 | male | left | Central America | 1,2,3 |
| MTLE 5 | 8 | 41 | 33 | 8 | 12 | female | right | Mexico | 1,2,3 |
| MTLE 6 | 15 | 40 | 25 | 18 | 6 | female | left | Mexico | 1,2,3 |
| MTLE 7 | 16 | 21 | 5 | 16 | 9 | female | left | Mexico | 1, 2 |
| MTLE 8 | 16 | 24 | 8 | 18 | 6 | female | right | Mexico | 1, 2 |
| MTLE 9 | 17 | 27 | 10 | 19 | 6 | male | right | Central America | 1,2,3 |
| MTLE 10 | 20 | 21 | 1 | Not Available | 11 | male | right | Central America | 1, 2 |
| MTLE 11 | 21 | 38 | 17 | 21 | 12 | female | left | Central America | 1,2,3 |
| MTLE 12 | 22 | 23 | 1 | 23 | 6 | male | right | Central America | 1, 2 |
| MTLE 13 | 22 | 31 | 9 | 23 | 12 | male | right | Central America | 1, 2 |
| MTLE 14 | 22 | 34 | 12 | 22 | 6 | male | left | Central America | 1, 2 |
| MTLE 15 | 23 | 35 | 12 | 26 | 5 | female | right | Mexico | 1, 2 |
| MTLE 16 | 25 | 37 | 12 | 26 | 5 | female | right | Mexico | 1, 2 |
| MTLE 17 | 25 | 32 | 7 | 25 | 5 | male | right | Mexico | 1, 2 |
| MTLE 18 | 26 | 38 | 12 | 27 | 6 | male | right | Mexico | 1,2,3 |
| MTLE 19 | 26 | 54 | 28 | 26 | 14 | female | right | Mexico | 1,2,3 |
| MTLE 20 | 27 | 27 | 0 | 31 | 12 | female | left | Mexico | 1, 2 |
| MTLE 21 | 27 | 39 | 12 | 30 | 1 | male | right | Mexico | 1,2,3 |
| MTLE 22 | 28 | 31 | 3 | 29 | 9 | male | left | Mexico | 1, 2 |
| MTLE 23 | 28 | 43 | 15 | 28 | 7 | female | left | Mexico | 1, 2 |
| MTLE 24 | 29 | 54 | 25 | Not Available | 14 | female | right | Mexico | 1, 2 |
| MTLE 25 | 31 | 39 | 8 | 31 | 6 | male | right | Mexico | 1,2,3 |
| MTLE 26 | 35 | 49 | 14 | 36 | 12 | female | right | Mexico | 1, 2 |
| MTLE 27 | 38 | 50 | 12 | 39 | 3 | female | right | Mexico | 1, 2 |
| MTLE 28 | 38 | 45 | 7 | 38 | 6 | female | right | Central America | 1,2,3 |
| MTLE 29 | 39 | 54 | 15 | 39 | 8 | female | left | Mexico | 1,2,3 |
| MTLE 30 | 42 | 48 | 6 | 42 | 8 | female | left | Mexico | 1,2,3 |
| MTLE 31 | 43 | 50 | 7 | 43 | 6 | male | left | Mexico | 1, 2 |
| MTLE 32 | 45 | 50 | 5 | 46 | 2 | female | left | Mexico | 1, 2 |
| MTLE 33 | 48 | 52 | 4 | Not Available | 6 | female | left | Central America | 1, 2 |
| MTLE 34 | 52 | 53 | 1 | 52 | 12 | male | left | Central America | 1,2,3 |
| MTLE 35 | 53 | 56 | 3 | 53 | 3 | female | right | Mexico | 1,2,3 |
| MTLE 36 | 56 | 57 | 1 | 56 | 12 | male | right | Mexico | 1,2,3 |

**Table S3:** Two-tailed Spearman’s Correlation between disease duration and presurgical cognitive z-scores. * P ≤ 0.05, ** P ≤ 0.01.

| Cognitive Domain | Number of patients | Spearman's correlation coefficient | p-value |
| --- | --- | --- | --- |
| Raven's | 34 | -0.4654 | 0.0055** |
| Digit Symbol | 36 | -0.2654 | 0.1178 |
| Color trails 1 | 35 | -0.2189 | 0.2064 |
| Color trails 2 | 34 | -0.2327 | 0.1853 |
| P-BNT | 36 | -0.1035 | 0.548 |
| COWAT/FAS | 33 | -0.1585 | 0.3783 |
| Digit Span | 36 | -0.2205 | 0.1963 |
| AVLT 5 | 36 | -0.4567 | 0.0051** |
| AVLT 7 | 36 | -0.3843 | 0.0207* |
| AVLT 8 | 36 | -0.254 | 0.135 |
| Block Design | 36 | -0.09061 | 0.5992 |
| Rey-O-CFT Copy | 36 | -0.4223 | 0.0103* |
| Rey-O-CFT Delay | 36 | -0.3401 | 0.0424* |
| Pegboard dominant | 34 | -0.2888 | 0.0976 |
| Pegboard Non-dominant | 34 | -0.314 | 0.0705 |

**Table S4:** A Critical period of cognitive impairment in MTLE patients. Three DD groups were created; G1: 1-20 years DD, G2: 21-30 years DD, and G3: 31+ years DD. Cognitive z-scores between 3 groups were compared with a one-way ANOVA with post-hoc Tukey’s analysis. *P ≤ 0.05, ** P ≤ 0.01.

| Cognitive Domain | F-statistic (DFn, DFd), p value | G1, G2, G3 (M, SD) | Tukey's multiple comparison test |
| --- | --- | --- | --- |
| **Raven's** | F (2, 31) = 5.097, p= 0.0122* | G1 (M = 0.062, SD = 0.70)<br>G2 (M = -0.85, SD = 0.98)<br>G3 (M = -0.89, SD = 0.56) | G1 vs G2 0.0266*<br>G1 vs G3 0.0195*<br>G2 vs G3 0.9897 |
| **Digit Symbol** | F (2, 33) = 2.141, p= 0.1336 | G1 (M = -0.23, SD = 0.86)<br>G2 (M = -1.1 SD = 1.1)<br>G3 (M = -0.90, SD= 0.93) | G1 vs G2 0.1273<br>G1 vs G3 0.2777<br>G2 vs G3 0.9109 |
| **Color trails 1** | F (2, 32) = 2.15, p= 0.1256 | G1 (M = 0.34, SD = 0.70)<br>G2 (M = -0.67 SD = 1.2)<br>G3 (M = -0.57, SD= 1.6) | G1 vs G2 0.1331<br>G1 vs G3 0.2259<br>G2 vs G3 0.9765 |
| **Color trails 2** | F (2, 32) = 1.691, p= 0.2004 | G1 (M = 0.016, SD = 0.84)<br>G2 (M = -0.1 SD = 1.7)<br>G3 (M = -0.79, SD= 0.43) | G1 vs G2 0.1870<br>G1 vs G3 0.3938<br>G2 vs G3 0.9107 |
| **P-BNT** | F (2, 33) = 0.4709, p= 0.6285 | G1 (M = -1.2, SD = 0.87)<br>G2 (M = -1.3 SD = 1.3)<br>G3 (M = -1.6, SD= 0.96) | G1 vs G2 0.9643<br>G1 vs G3 0.6332<br>G2 vs G3 0.7485 |
| **COWAT/FAS** | F (2, 30) = 0.4559, p= 0.6382 | G1 (M = -0.28, SD = 0.90)<br>G2 (M = -0.45 SD = 1.0)<br>G3 (M = -0.70, SD= 0.99) | G1 vs G2 0.9190<br>G1 vs G3 0.6204<br>G2 vs G3 0.8131 |
| **Digit Span** | F (2, 33) = 1.994, p= 0.1522 | G1 (M = -0.32, SD = 1.3)<br>G2 (M = 0.12 SD = 1.3)<br>G3 (M = -0.93, SD= 1.4) | G1 vs G2 0.7123<br>G1 vs G3 0.5391<br>G2 vs G3 0.1293 |
| **AVLT 5** | F (2, 33) = 7.096, p = 0.0027** | G1 (M = 0.18, SD = 0.48)<br>G2 (M = -1.0, SD= 1.1)<br>G3 (M = -1.2, SD = 0.95) | G1 vs G2 0.0094**<br>G1 vs G3 0.0040**<br>G2 vs G3 0.8853 |
| **AVLT 7** | F (2, 33) = 5.874, p= 0.0066 ** | G1 (M = -0.13, SD = 1.2)<br>G2 (M = -1.5 SD = 0.99)<br>G3 (M = -1.6, SD= 1.1) | G1 vs G2 0.0146*<br>G1 vs G3 0.0112*<br>G2 vs G3 0.9736 |
| **AVLT 8** | F (2, 33) = 3.906, p= 0.0300* | G1 (M = -0.43, SD = 1.3)<br>G2 (M = -1.7 SD = 0.96)<br>G3 (M = -1.6, SD= 1.3) | G1 vs G2 0.0373*<br>G1 vs G3 0.0645<br>G2 vs G3 0.9832 |
| **Block Design** | F (2, 33) = 1.200, p= 0.3141 | G1 (M = -0.14, SD = 1.2)<br>G2 (M = -0.82, SD = 1.0)<br>G3 (M = -0.62, SD= 1.0) | G1 vs G2 0.2878<br>G1 vs G3 0.5609<br>G2 vs G3 0.8764 |
| **Rey-O-CFT Copy** | F (2, 33) = 2.842, p= 0.0727 | G1 (M = 0.53, SD = 0.58)<br>G2 (M = -0.22, SD = 1.4)<br>G3 (M = -0.56, SD= 0.91) | G1 vs G2 0.2341<br>G1 vs G3 0.0629<br>G2 vs G3 0.7027 |
| **Rey-O-CFT Delay** | F (2, 33) = 1.622, p= 0.2129 | G1 (M = -0.065, SD = 1.1)<br>G2 (M = -0.69, SD = 1.3)<br>G3 (M = -0.91, SD= 0.82) | G1 vs G2 0.3807<br>G1 vs G3 0.2032<br>G2 vs G3 0.8786 |
| **Pegboard dominant** | F (2, 31) = 5.219, p= 0.011* | G1 (M = -0.65, SD = 1.2)<br>G2 (M = -2.3 SD = 1.3)<br>G3 (M = -2.0, SD= 1.2) | G1 vs G2 0.0108*<br>G1 vs G3 0.0547<br>G2 vs G3 0.8134 |
| **Pegboard Non-dominant** | F (2, 31) = 3.938, p= 0.0299* | G1 (M = -0.98, SD = 1.3)<br>G2 (M = -2.3 SD = 1.3)<br>G3 (M = -2.1, SD= 0.93) | G1 vs G2 0.0333*<br>G1 vs G3 0.0815<br>G2 vs G3 0.9409 |

**Table S5:** Two-tailed Spearman’s Correlation between presurgical cognitive z-scores and the number of 1) immature Dcx+ Prox1+ cells/mm^3^ and 2) mature Prox1+ cells/mm^3^ detected from the surgically resected hippocampus. * P ≤ 0.05.

| Cognitive Domain | Number of patients | Immature Neurons (Dcx+ Prox1+) |  | Mature Neurons (Prox1+) |  |
| --- | --- | --- | --- | --- | --- |
|  |  | Spearman's correlation coefficient (r) | p-value | Spearman's correlation coefficient (r) | p-value |
| Raven's | 15 | 0.3819 | 0.159 | -0.06077 | 0.8296 |
| Digit Symbol | 17 | 0.1476 | 0.5681 | 0.114 | 0.6611 |
| Color trails 1 | 17 | 0.3274 | 0.1977 | 0.26 | 0.3111 |
| Color trails 2 | 16 | 0.2438 | 0.3608 | 0.5457 | 0.0311* |
| P-BNT | 17 | 0.2796 | 0.274 | 0.2374 | 0.356 |
| COWAT/FAS | 14 | 0.5833 | 0.0315* | 0.5969 | 0.0268* |
| Digit Span | 17 | 0.2274 | 0.3761 | 0.1034 | 0.6913 |
| AVLT 5 | 17 | 0.4969 | 0.0443* | 0.3229 | 0.205 |
| AVLT 7 | 17 | 0.2537 | 0.3226 | 0.02217 | 0.9356 |
| AVLT 8 | 17 | 0.3707 | 0.1423 | 0.003727 | 0.9904 |
| Block Design | 17 | 0.2253 | 0.3806 | -0.3227 | 0.2053 |
| Rey-O-CFT Copy | 17 | 0.2766 | 0.2793 | -0.05763 | 0.826 |
| Rey-O-CFT Delay | 17 | -0.02195 | 0.9345 | -0.2083 | 0.4208 |
| Pegboard dominant | 15 | 0.4147 | 0.1248 | 0.1117 | 0.6898 |
| Pegboard Non-dominant | 15 | 0.3842 | 0.1569 | 0.3802 | 0.1617 |

**Figure S1.**
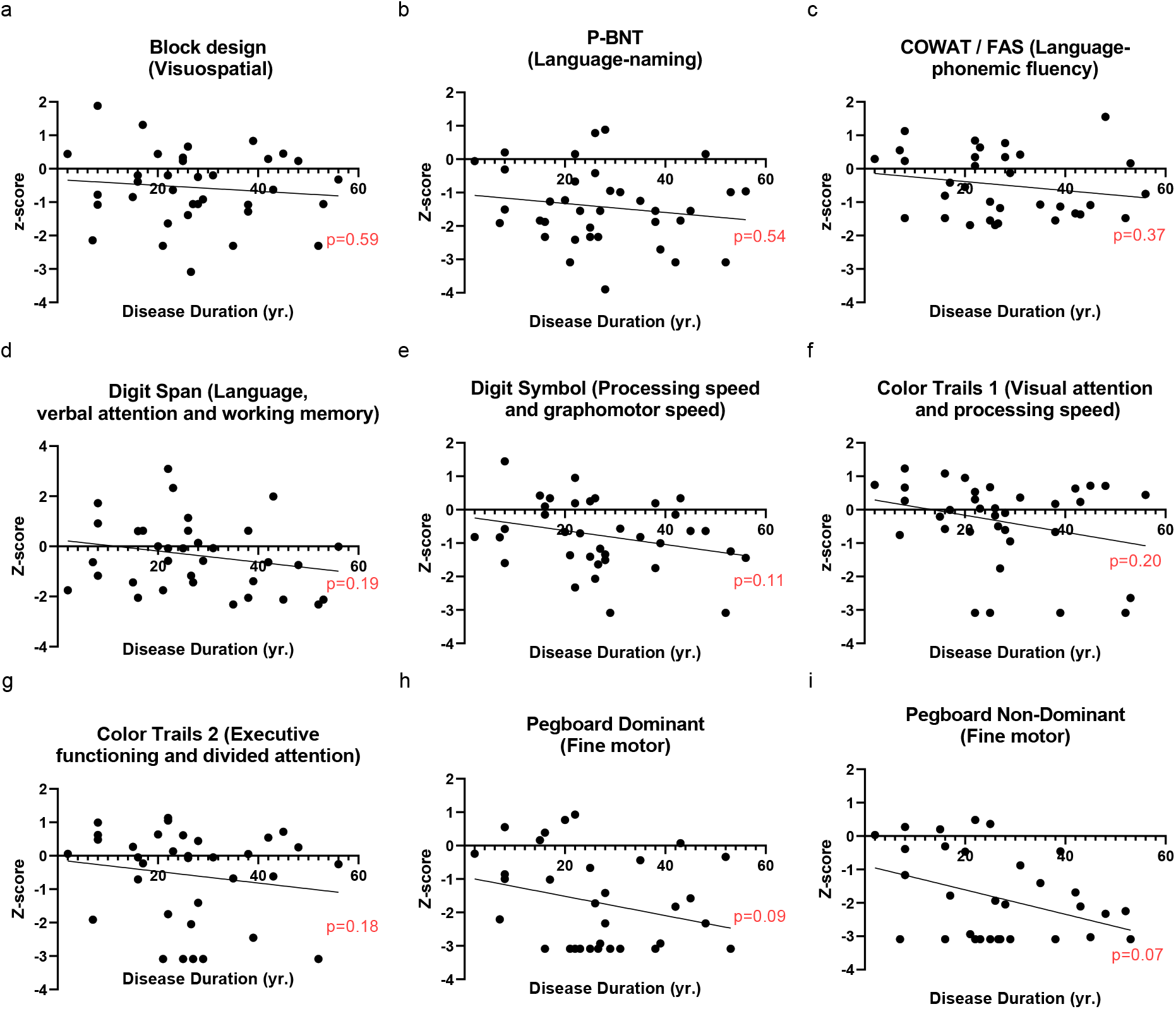
Cognitive domains that do not have a significant decline over disease duration (DD). Association between DD and cognitive z scores calculated by two-tailed Spearman’s correlation for NeSBHIS test: a) Block Design visuospatial, b) P-BNT language-naming, c) COWAT/FAS language fluency, d) Digit Span language attention, e) Digit Symbol processing speed, f) Color trails 1 visual attention, g) Color trails 2 executive functioning, h) Pegboard dominant fine motor control, i) Pegboard Non-dominant fine motor control.

**Figure S2:**
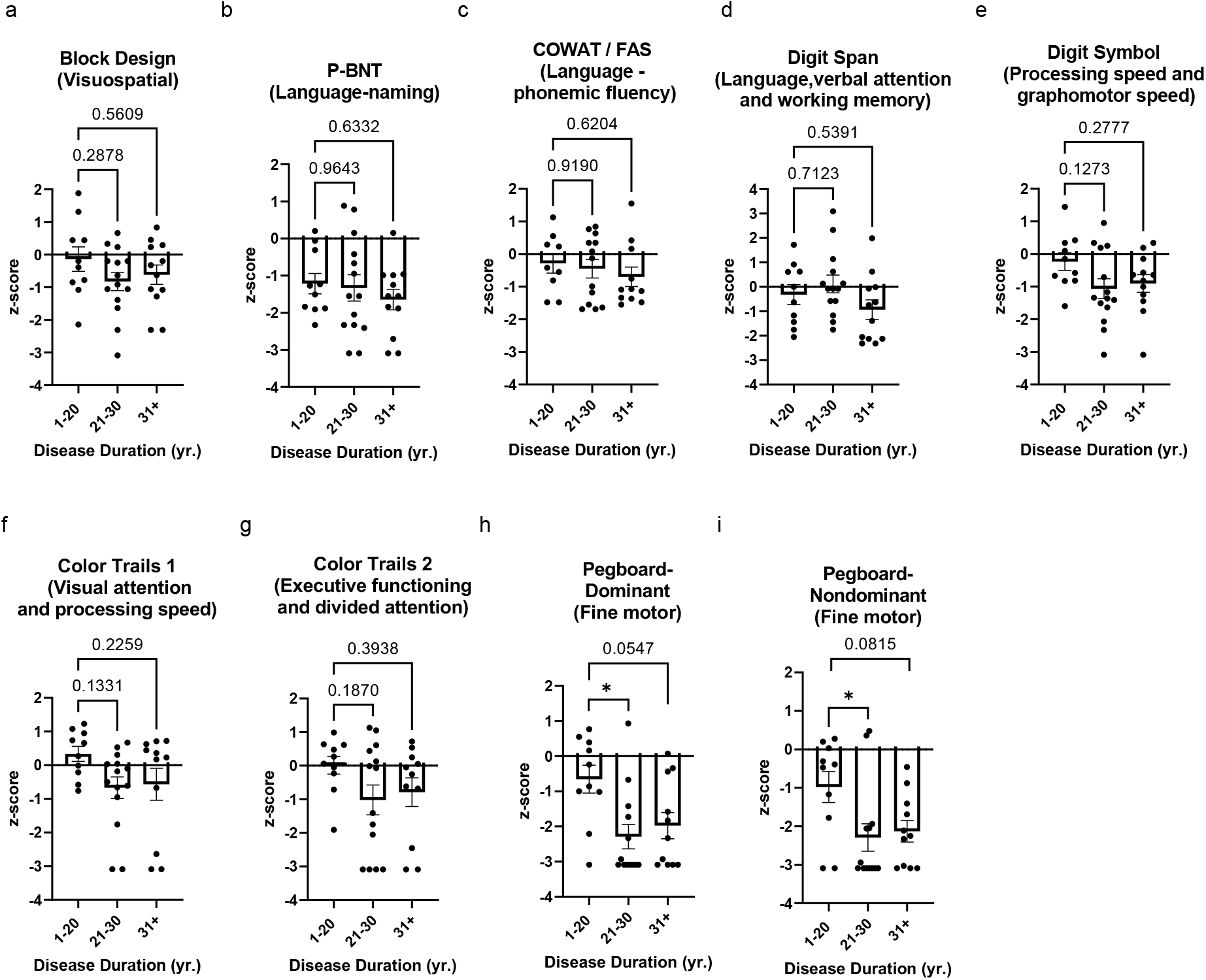
Cognitive domains which did not drop in performance at a critical 20 year disease duration (DD). Three DD groups were created; G1: 1-20 years DD, G2: 21-30 years DD, and G3: 31+ years DD. ANOVA (post-hoc Tukey’s) analysis was performed between the three groups. a-g) Cognitive z scores were not significantly different between groups. a) Block design visuospatial, b) P-BNT language-naming, c) COWAT/FAS language fluency, d) Digit span language attention, e) Digit symbol processing speed, f) Color trails 1 visual attention, g) Color trails 2 executive functioning. h, i) Pegboard-Dominant and Non-dominant fine motor control showed significant difference between groups with G1 significantly higher than G2, but not compared to G3. *P ≤ 0.05. All error bars represent S.E.M across individual cases.

**Figure S3:**
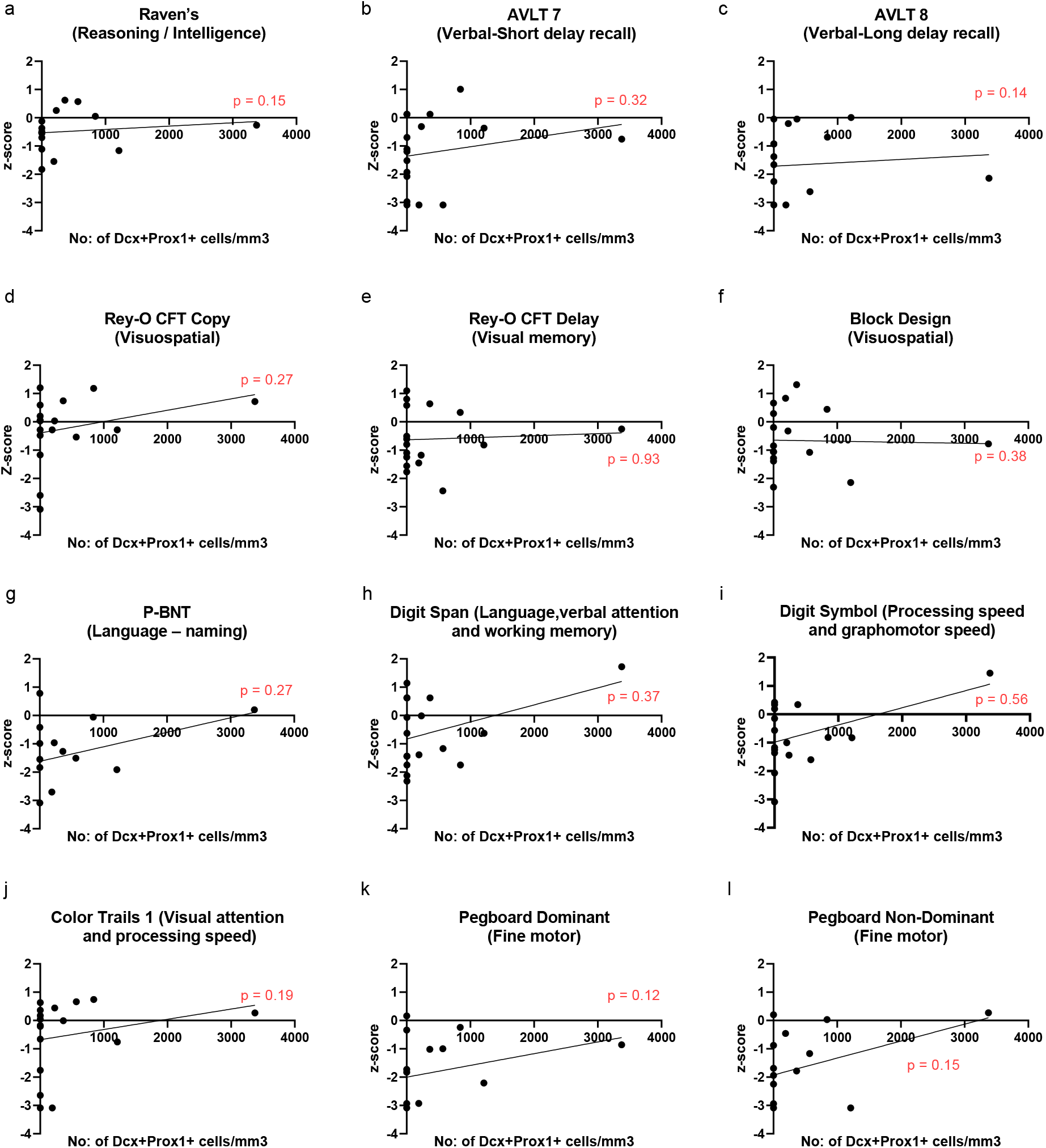
Cognitive domains lacking significant association with neurogenesis levels. Two-tailed Spearman’s Correlation between the number of Dcx+ Prox1+ immature neurons/mm3 detected from the surgically resected hippocampus and presurgical cognitive z-scores for a) Raven’s Intelligence, b) AVLT 7 short-delay recall, c) AVLT 8 long-delay recall, d) Rey-O-CFT Copy visuospatial, e) Rey-O-CFT Delay visuospatial memory, f) Block Design visuospatial, g) P-BNT language, h) Digit Span language attention, i) Digit Symbol processing speed, j) Color trails 1 visual attention, k) Pegboard dominant fine motor control, l) Pegboard Non-dominant fine motor control.

**Figure S4:**
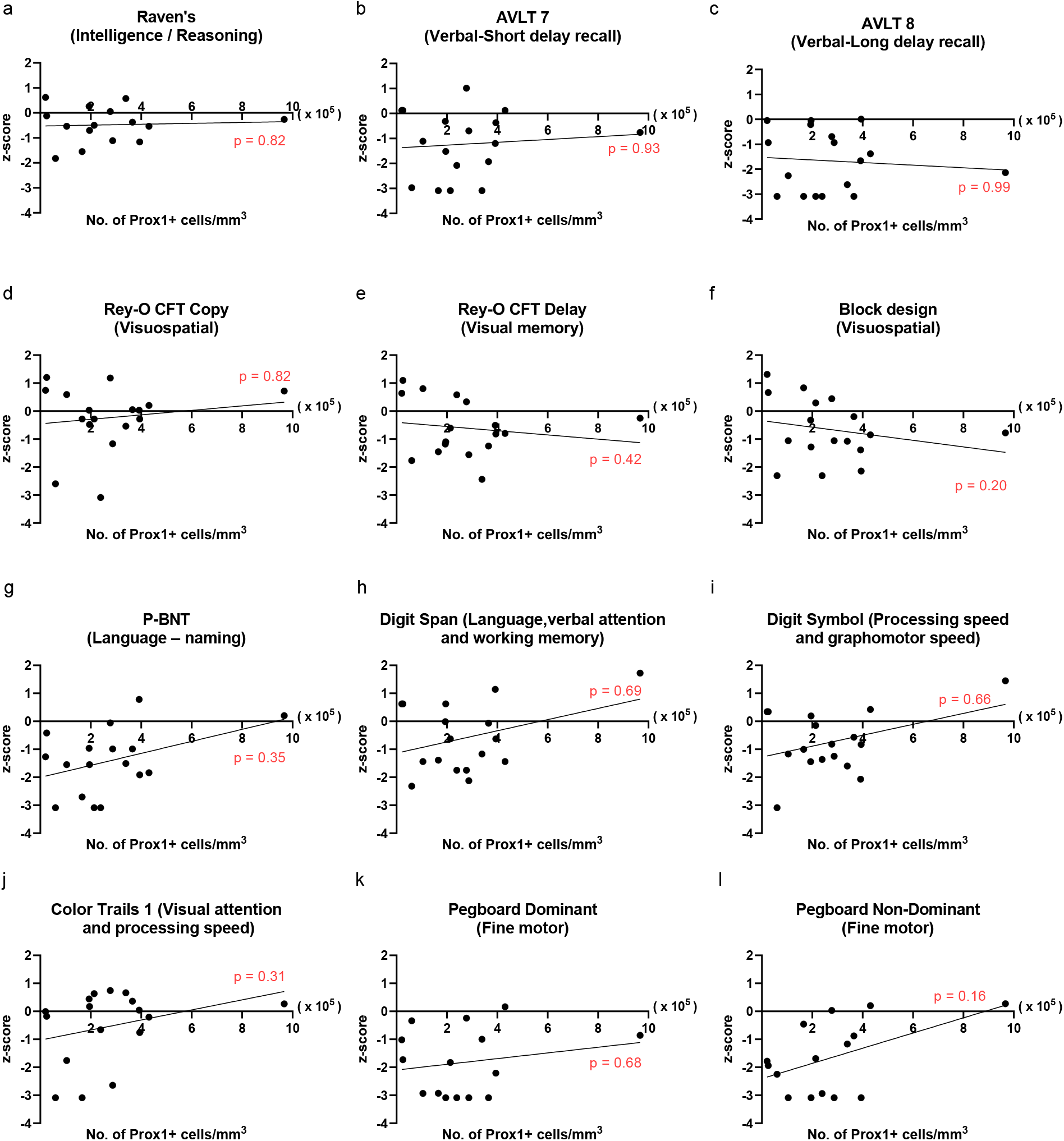
Cognitive domains lacking significant association with mature granule neuron levels. Two-tailed Spearman’s Correlation between the number of Prox1+ granule neurons/mm3 detected from the surgically resected hippocampus and presurgical cognitive z-scores for a) Raven’s Intelligence, b) AVLT 7 short-delay recall, c) AVLT 8 long-delay recall, d) Rey-O-CFT Copy visuospatial, e) Rey-O-CFT Delay visuospatial memory, f) Block Design visuospatial, g) P-BNT language, h) Digit Span language attention, i) Digit Symbol processing speed, j) Color trails 1 visual attention, k) Pegboard dominant fine motor control, l) Pegboard Non-dominant fine motor control.

## ONLINE METHODS

### Cognitive assessment

Retrospective cognitive analysis of presurgical neuropsychological exams of Spanish-speaking patients (N=36) with intractable TLE was performed to explore the relationship between DD and TLE with cognitive functioning. The study was approved by University of Southern California Institutional Review Board approval number: HS-10-00162, and Rancho Los Amigos National Rehabilitation Center and the Rancho Research Institute’s Institutional (RRI) Review Board, approval number: RRI IRB # 195 and the data use agreement with RRI and the University of Texas Southwestern Medical Center IRB Protocol approval number: STU-052018-086.

We have complied with all relevant ethical regulations. Written consents were obtained from the patients to study the clinical and demographic information. De-identified patient clinical details (age and DD of epilepsy at the time of Neuropsychological test, age of disease onset, sex, left vs right hemisphere of effected hippocampus, education years, ethnicity, nationality) are provided in Extended Data Table 2. Both male (N = 16) and female (N = 20) cases were included to examine sex as a biological variable in the study. The NeSBHIS was part of a comprehensive presurgical evaluation. Intellect, attention, processing speed, language, visuospatial, visual memory, verbal learning and memory, executive functioning and fine motor dexterity were measured to assess epilepsy comorbidities (Table S1). To account for variability in education, which influences neuropsychological test performance^22^, subtest raw scores were converted to z-scores using education-based normative data^46^. Grooved pegboard was used instead of the fine-motor Pin test from NeSBHIS, which was normalized using age and education based normative data as in Smith, et.al, 2020^22^. Upper and lower cut off values for z scores were set at +/- 3.09 to eliminate potential outliers due to very superior performance (z scores > +3.09) and clinically profound impairment (<3.09).

### Human MTLE Hippocampal Tissue processing

Human hippocampal tissue was obtained from 17 out of 36 patients undergoing an *en bloc* resection of their hippocampus for the treatment of medically resistant MTLE, following strict ethical guidelines. Written consents for the surgical treatment of MTLE, tissue donation for the study as well as relevant clinical and demographic information collection (University of Southern California Institutional Review Board approval number: HS-17-00370) were obtained from all the patients prior to the surgery. Both male (N = 8) and female (N = 9) cases were included to examine sex as a biological variable in the study. The surgery was performed by neurosurgeons at the Keck Medical Center or Los Angeles County Hospital, University of Southern California. The surgery was performed within a year of neuropsychology test for 10 out of 17 patients. The rest of the cases underwent surgery within 4 years of test (N = 4 after 1 year, N = 1 after 2 years, N = 2 after 3 years). DD at surgery is provided in Extended Data Table 2. Surgically resected hippocampal tissue was transported to the laboratory within 30 minutes to 1 hour of surgical removal in a tube containing HypoThermosol^®^ FRS (BioLife Solutions, Bothell, WA) solution maintained at approximately 4°C in an ice box.

### Tissue Processing

Hippocampal tissue was cut into 400 µm thick slices in chilled (4°C), oxygenated (95% O_2_, 5% CO_2_) N-methyl-D-glycamine - artificial cerebrospinal fluid (aCSF) solution (NMDG-aCSF; NMDG 93mM; KCl 2.5mM; NaH_2_PO_4_ 1.2mM; NaHCO_3_ 30mM; HEPES 20mM; Glucose 25mM; Sodium ascorbate 5mM; Thiourea 2mM; Sodium pyruvate 3mM; MgSO_4_ 10mM; CaCl_2_ 0.5mM) using a vibrating microtome (Leica VT1200S, Leica Biosystems, Germany). Vertical deflection of the blade was minimized with Vibrocheck technology (Leica Biosystems, Germany). Slicing parameters were set at speed 0.5-0.15 mm/s and vibration amplitude 1.5 mm. Hippocampal slices were first used for multi electrode array experiments explained in our previous study^18^. Slices were allowed to recover in a chamber with warm (32°C) aCSF (NaCl 124 mM; KCl 4 mM; NaHCO_3_ 26 mM; Glucose 10 mM; CaCl_2_ 2mM; MgCl_2_ 2mM) solution (30°C, 95% O_2_, 5% CO_2_) for 1 hour before multi-electrode array (MEA) recording. Continuous oxygenated, warm (32-34^◦^C) aCSF at a rate of 6 ml/min was circulated in the MEA ring well during tissue recording. A 5 min baseline recording was performed in aCSF followed by a 20-30 min recording in a modified aCSF solution containing high (8 mM) potassium, low (0.25 mM) magnesium and 100 µM 4-aminopyridine (4- AP) to induce inter-ictal like activity. Detailed MEA method is explained in our previous study identifying neurogenesis in human MTLE^18^. Following the MEA recording, slices were immediately fixed using 4% PFA for immunohistochemistry.

### Immunohistochemistry and cell quantification

Slices were pre-treated in a similar manner for methodological consistency between cases. Slices were fixed in 4% PFA - made freshly from 16% PFA stored at -20°C - for 30 minutes at room temperature and washed thoroughly with 1x phosphate buffered saline (PBS) overnight. The slices were dehydrated using 30% sucrose in PBS at 4 °C. After 1 week, once the slice sank it was embedded in O.C.T compound (Tissue-Tek), ensuring the DG region is flat. Slices were further sub-sectioned by cryostat (Leica CM 3050 S) into 30µm and mounted to super frost plus slides. The slides were dried overnight at room temperature and transferred to -20°C for long term storage. Before performing immunohistochemistry, slides were equilibrated to room temperature for 2 hours or at 37°C for 30 minutes. Slides were further washed in PBS for 1 hour, followed by blocking and permeabilization in 3% Bovine serum albumin (BSA), 10% Donkey serum in 0.1% Triton-X in PBS for 2 hours at room temperature. The sections were incubated with primary antibodies for 18-20 hours at 4°C in 0.3 % BSA, 1% Donkey serum and 0.1% Triton X in PBS. Following a 1-hour equilibration to room temperature, slides were washed in 0.1% Triton X in PBS for 15 minutes 3 times. Secondary antibodies from Jackson Immuno research laboratories were diluted from a stock solution of 0.6mg/ml and incubated along with 1µg/ml of DAPI (Roche, Cat No: 10236276001) for 2 hours at room temperature in 0.3 % BSA, 1% Donkey serum and 0.1% Triton X. Sections were washed 3 times in 0.1% Triton X in PBS for 15 minutes and mounted with cover glass. The whole DG region was imaged at 20X using a Zeiss LSM 700 confocal microscope automated tile scan. All cell quantification was performed manually in Zeiss blue software. GCL boundaries were marked using the DAPI channel around the seemingly compact cell layer leaving a gap of two cell nuclear distance on both sides of the hilus and molecular layer. Each whole section was analysed to quantify the number of rare Dcx+ Prox1+ cells. Representative sub regions in the GCL was analysed for mature Prox1+ granule neurons due to their high prevalence. The remaining statistical analysis and plots were prepared using Prism8 from GraphPad Inc.

### List of antibodies

| **Primary Antibodies** |  |  |  |  |
| --- | --- | --- | --- | --- |
| **Antibodies** | **Species** | **Dilution** | **Company** | **Catalog No:** |
| Doublecortin (Dcx) | Mouse | 3 in 100 | Santa Cruz | sc-390645 |
| Prox1 | Rabbit | 1 in 200 | Abcam | ab101851 |
| **Secondary Antibodies** |  |  |  |  |
| **Antibodies** | **Species** | **Dilution** | **Company** | **Catalog No:** |
| Anti-mouse Alexa Fluor 488 | Donkey | 1 in 200 | Jackson ImmunoResearch | 715-546-151 |
| Anti-rabbit Alexa Fluor 488 | Donkey | 1 in 200 | Jackson ImmunoResearch | 711-546-152 |
| Anti-mouse Cy3 | Donkey | 1 in 200 | Jackson ImmunoResearch | 715-166-150 |
| Anti-rabbit Cy3 | Donkey | 1 in 200 | Jackson ImmunoResearch | 711-166-152 |
| Anti-rabbit Alexa Fluor 647 | Donkey | 1 in 200 | Jackson ImmunoResearch | 711-606-152 |

### Statistical Analysis

All the statistical analysis was performed using GraphPad Prism 8. The statistical methods used for each Figure is specifically noted in the Figure legends. A two tailed Spearman’s correlation was performed to calculate the strength of linear relationship between two variables (Figure 1 and 3). For Figure 2 to identify the critical window of cognitive decline, the cohort was divided into three DD groups; < 20 years (G1, N = 10), 21 to 30 years (G2, N =14), and > 31-years (G3, N =12), allowing ANOVA analyses, followed by a Tukey’s multiple comparison post-hoc test. For Figure 3c, a comparison of fits between linear regression, logarithmic, and exponential one phase decay was performed. The parameters for goodness of fit were better (R^2^ value closer to 1, lower RMSE) for exponential compared to logarithmic and linear regressions.

| | $R^2$ | RMSE |
| --- | --- | --- |
| Linear | 0.28 | 714.3 |
| Exponential one phase decay | 0.41 | 646.2 |
| Logarithmic | 0.35 | 704.66 |

